# *Pseudodesulfovibrio cashew* sp. nov., a novel deep-sea sulfate-reducing bacterium, linking heavy metal resistance and sulfur cycle

**DOI:** 10.1101/2020.12.08.417204

**Authors:** Rikuan Zheng, Chaomin Sun

**Affiliations:** Key Laboratory of Experimental Marine Biology, Institute of Oceanology, Chinese Academy of Sciences, Qingdao 266071, China; Laboratory for Marine Biology and Biotechnology, Qingdao National Laboratory for Marine Science and Technology, Qingdao 266071, China; College of Earth Science, University of Chinese Academy of Sciences, Beijing 100049, China; Center of Ocean Mega-Science, Chinese Academy of Sciences, Qingdao, 266071, China

**Keywords:** dissimilatory sulfate reduction, sulfur cycle, heavy metals, *dsr* genes, deep sea, cold seep

## Abstract

Sulfur cycling is primarily driven by sulfate reduction mediated by sulfate-reducing bacteria (SRB) in marine sediments. The dissimilatory sulfate reduction drives the production of enormous quantities of reduced sulfide and thereby the formation of highly insoluble metal sulfides in marine sediments. Here, a novel sulfate-reducing bacterium designated *Pseudodesulfovibrio cashew* SRB007 was isolated and purified from the deep-sea cold seep and proposed to represent a novel species in the genus of *Pseudodesulfovibrio.* A detailed description of the phenotypic traits, phylogenetic status and central metabolisms of strain SRB007, allowing the reconstruction of the metabolic potential and lifestyle of a novel member of deep-sea SRB. Notably, *P. cashew* SRB007 showed a strong ability to resist and remove different heavy metal ions including Fe^3+^, Co^2+^, Ni^2+^, Cu^2+^, Cd^2+^ and Hg^2+^. And the dissimilatory sulfite reduction was demonstrated to contribute to the prominent removal capability of *P. cashew* SRB007 against different heavy metals via forming insoluble metal sulfides.

**IMPORTANCE:** The dissimilatory sulfate reduction driven by sulfate-reducing bacteria (SRB) was ubiquitous in marine sediments, and was proposed to couple with heavy metal ions removal through forming insoluble metal sulfides. The deep-sea cold seep is a very special environment where is rich in sulfate and novel species of SRB that possessing many unknown mechanisms toward sulfur cycle. Here, a novel sulfate-reduction bacterium *Pseudodesulfovibrio cashew* SRB007 was isolated from the deep-sea cold seep and proposed as the type strain for a novel species. The taxonomy and typical physiological properties closely related to sulfur cycle, heavy metal resistance and their co-relationship were disclosed through a combination of genomic and biochemical methods. Given the absence of pure cultures of typical SRB isolated from the deep-sea cold seep, our work provides a good model to study the sulfur cycle which coupling with other elements and a potential candidate to develop bioremediation product in the future.

## INTRODUCTION

Sulfur is an essential element for life, which is widely found in the natural environment. The ocean represents a major reservoir of sulfur on Earth, with large quantities in the form of dissolved sulfate ion (SO_4_^2-^), which is the second most abundant anion next to chloride (1). Marine sediments are the main sink for sea-water sulfate and the sedimentary sulfur cycle is a major component of the global sulfur cycle (2, 3). It is noting that the sulfur cycle of marine sediments is primarily driven by the dissimilatory sulfate reduction (DSR), which is mediated by sulfate-reducing bacteria (SRB) in many anaerobic environments (2). SRB mediate two sulfate reduction pathways: assimilative sulfate reduction (ASR) and dissimilating sulfate reduction (DSR). Ecologically, DSR plays a major role in the global sulfur and carbon cycle (4). The canonical microbial pathway for dissimilatory sulfate reduction involves the initial reduction of sulfate (SO_4_^2-^) to sulfite (SO_3_^2-^) by a combination of sulfate adenylyltransferase (Sat) and adenylyl-sulfate reductase (AprAB), followed by reduction of SO_3_^2-^ to H_2_S (S^2-^) by two different pathways. One is that SO_3_^2-^ is reduced to H_2_S in a single step by a single enzyme, bisulfite reductase (5). And the other pathway involves several enzymes (such as sulfite reductase, trithionate reductase and thiosulfate reductase) and intermediates (such as trithionates and thiosulfates) (6).

Given that the formation of a large amount of sulfide in the course of sulfate reduction and most of the heavy metals react with sulfide to form highly insoluble metal sulfides, SRB-mediated dissimilating sulfate reduction is proposed an effective way to cope with the stress of many harmful metal ions which broad distribute in marine sediments (7, 8). Meanwhile, an accepted clever idea was developed by researchers to remove heavy metals from the environment by utilizing the dissimilating sulfate reduction mediated by different SRB (9, 10). Hence, SRB-mediated metal sulfide precipitation represents a potentially useful way for the bioremediation of metal ion contaminants, which was an attractive alternative over physicochemical methods (9, 11).

Overall, rather than being a simple cycle, composed of anaerobic bacterial reduction of sulfate to hydrogen sulfide and aerobic reoxidation of H_2_S to SO_4_^2-^, the transformations of sulfur in marine sediments also links to the cycles of carbon, nitrogen, iron, manganese and other important elements (12–14). Therefore, it is of utmost importance to identify novel SRB mediating sulfur cycle in marine sediments, which will be of great advantage to disclose novel mechanisms and develop more powerful bioremediation products. *Pseudodesulfovibrio* is a new genus of SRB, which was originally proposed and reclassified of four species of the genus *Desulfovibrio* by Cao *et al* in 2016 (15). Till to date, most species of the genus *Pseudodesulfovibrio* were isolated from marine sediments, including *Pseudodesulfovibrio profundus* (16), *Pseudodesulfovibrio portus* (17), *Pseudodesulfovibrio piezophilus* (18) and *Pseudodesulfovibrio indicus* (15), which strongly indates that this novel genus play important roles in driving the sulfur cycle in marine sediments. However, till to date, there are no any results about the sulfur cycle and heavy metal resistance mediated by *Pseudodesulfovibrio* other than taxonomic data have been published.

In the present study, a novel species of the genus *Pseudodesulfovibrio*, SRB007, was isolated from the deep-sea cold seep and proposed as the type strain for this novel species. Furthermore, the taxonomy and typical physiological properties closely related to sulfur cycle and heavy metal resistance were disclosed through the combination of genomic and biochemical methods, providing a clue to understand the coupling of different elements in the deep-sea cold seep and a potential candidate to develop bioremediation product in the future.

## RESULTS

### Isolation and identification of a novel deep-sea sulfate-reducing bacterium *P. cashew* SRB007

During isolation of uncultured microorganisms from the deep-sea cold seep, a potential novel species (strain SRB007) of SRB was obtained after several rounds of purification, which showing 97.34% similarity of 16S rRNA sequence to that of *Pseudodesulfovibrio profundus* DSM 11384^T^, the type strain of the genus *Pseudodesulfovibrio*, isolated from a deep sediment layer in the Japan Sea (16). Strain SRB007 was mesophilic, strictly anaerobic and Gram-stain-negative, while spores were never observed. Under transmission electron microscopy (TEM) observation, the cells of strain SRB007 were cashew-shaped, approximately 1.0-2.5 × 0.3-0.7 μm in size and had peritrichous flagella (Figs. 1A and 1B). To further identify the taxonomic status of strain SRB007, we further performed the phylogenetic analyses with 16S rRNA genes from some cultured representatives of the family *Desulfovibrionaceae*. All of the phylogenetic trees of 16S rRNA showed that strain SRB007 fell within the cluster comprising *Pseudodesulfovibrio* species and the closest species in the NCBI database was *Pseudodesulfovibrio profundus* DSM 11384^T^ (97.34% sequence similarity) (16), and the next more closely related recognized species were *Pseudodesulfovibrio piezophilus* C1TLV30^T^ (96.68% similarity) (18) and *Pseudodesulfovibrio indicus* J2^T^ (95.71% similarity) (Figs. 1C, Fig. S1 and Fig. S2).

**FIG 1.**
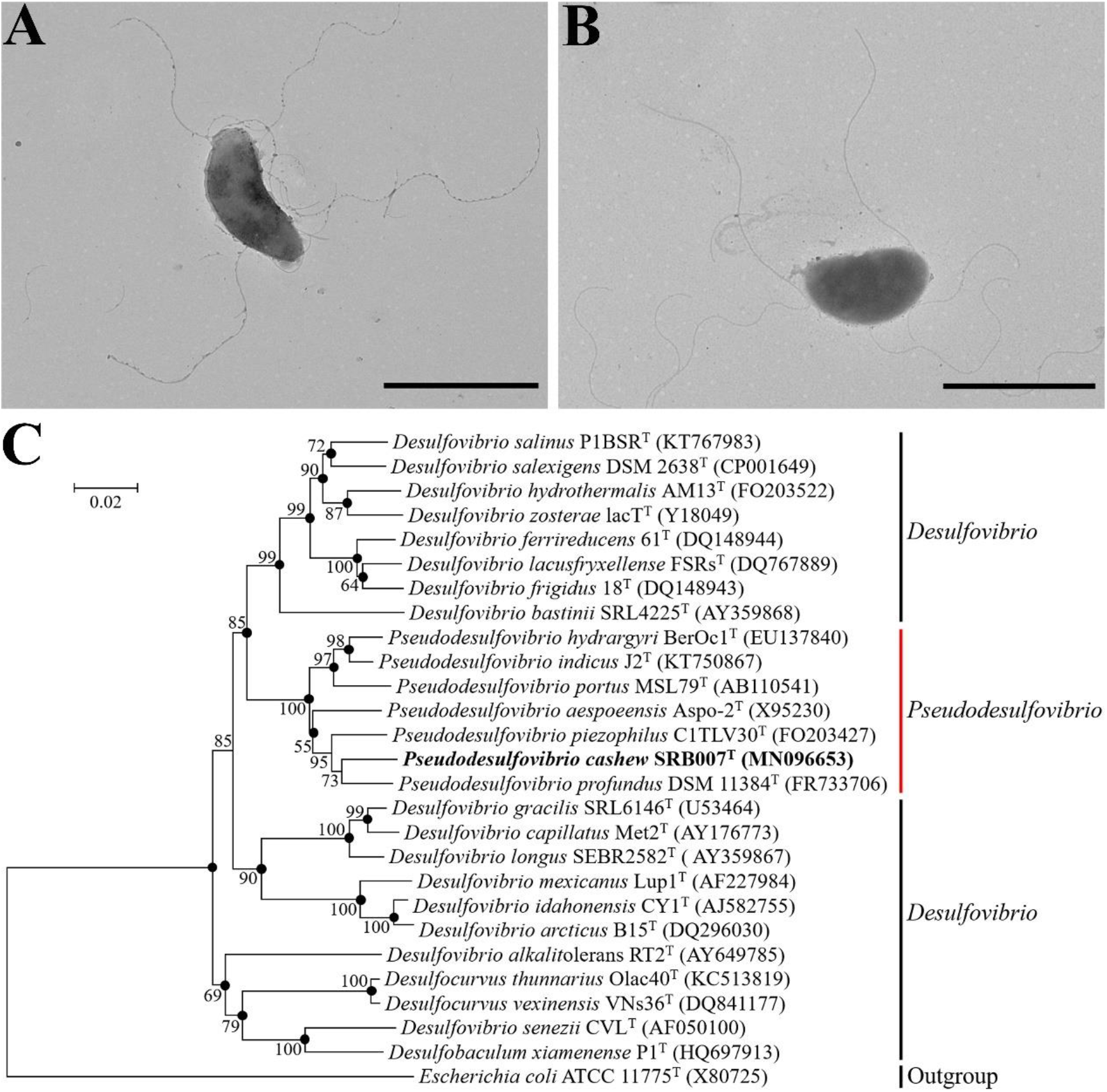
Morphology observation and phylogenetic analysis of *P. cashew* SRB007. (A and B) Morphology observation of *P. cashew* SRB007 by TEM. Bars, 2 μm. (C) Phylogenetic tree based on 16S rRNA gene sequences of *P. cashew* SRB007 and related strains reconstructed by the neighbor-joining method. GenBank accession numbers are given in parentheses after strain names. Bootstrap values are based on 1,000 replicates. Bar, 0.02 substitution per nucleotide position. *Escherichia coli* ATCC 11775^T^ (GenBank accession number X80725) was used as the outgroup.

To further confirm the novel species position of *P. cashew* SRB007, the dissimilatory sulfite reductases *(dsrAB)* sequences of *P. cashew* SRB007 and the most closely related species belonging to the general *Pseudodesulfovibrio*, *Desulfovibrio* and *Desulfomicrobium*, were extracted from the corresponding genome and then applied for phylogenetic analysis (Table S1). Consistent with the result of 16S rRNA-based phylogenetic analysis, the result of *dsrAB*-based phylogenetic tree also placed *P. cashew* SRB007 as a sister clade of *P*. *profundus* DSM 11384^T^ within the genus *Pseudodesulfovibrio* (Fig. S3). A sequence similarity calculation using the NCBI server indicated that the closest relatives of strain SRB007 were *P. profundus* DSM 11384^T^ (88.32% similarity), *P. aespoeensis* Aspo-2^T^ (86.78% similarity), *P. indicus* J2^T^ (86.40% similarity) and *P. piezophilus* C1TLV30^T^ (85.19% similarity). These results allowed us to propose the strain SRB007 as a representative of a novel species belonging to the *Pseudodesulfovibrio* genus. Thus, the strain SRB007 was proposed as the type strain and designated as *Pseudodesulfovibrio cashew* SRB007^T^.

### Physiological and chemotaxonomic characteristics of *P. cashew* SRB007

To further gain insights into the lifestyle of *P. cashew* SRB007, a series of physiological and chemotaxonomic characteristics of this bacterium were investigated. *P. cashew* SRB007 had a high survival ability to tolerate different salt concentrations (0-100 g/L NaCl) (Table 1), which was according well with other typical SRB isolated from the ocean (18–24) (Supplementary Table S2), and this property might benefit this bacterium to survive in various hypersaline habitats. Compared with the closely related type strain *P. profundus* DSM 11384^T^ and other SRB,*P. cashew* SRB007 showed a wider range to utilize different substrates as electron donors (including acetate, fumarate, malate, succinate, formate, lactate, methanol and ethanol) and acceptors (including sulfate, sulfite, thiosulfate, nitrate and nitrite) (Table 1), which endowed *P. cashew* SRB007 with a strong flexibility to different environments.

**Table 1.**
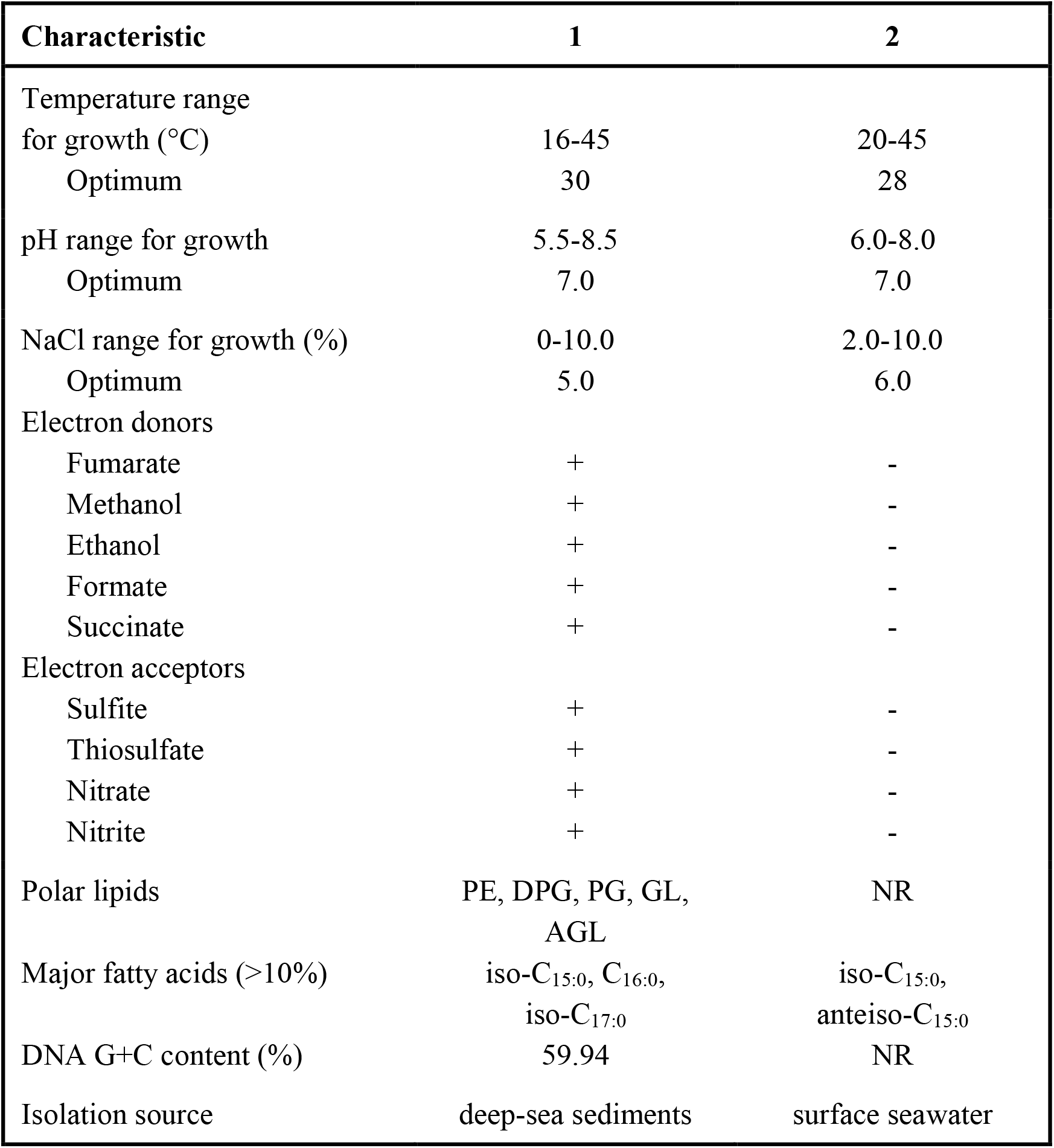
Differential physiological characteristics of *P. cashew* SRB007^T^ and the closely related type strain *P. profundus* DSM 11384^T^. Strains: 1, *P. cashew* SRB007^T^(all data from this study); 2, *P. profundus* DSM 11384^T^ (all data from this study except DNA G+C content and polar lipids). **+**, Positive result or growth; -, negative result or no growth; NR, not reported.

The major polar lipids in *P. cashew* SRB007 were phosphatidylethanolamine, diphosphatidylglycerol, phosphatidylglycerol, unidentified glycolipid and unknown aminoglycolipids (Fig. S5). The predominant fatty acids (>10%) were iso-C_15:0_, C_16:0_ and iso-C_17:0_ (Table 1). The amount of iso-C_15:0_, C_16:0_ and iso-C_17:0_ in *P. cashew* SRB007 (38.87%, 21.28% and 11.10%) were higher than those found in *P. profundus* DSM 11384^T^ (22.61%, 7.50% and 1.15%, respectively), while the amount of anteiso-C_15:0_ in *P. cashew* SRB007 (8.80%) was lower than that found in *P. profundus* DSM 11384^T^ (15.22%) (Table 2). The distinctive composition of fatty acids of *P. cashew* SRB007 facilitates it to better adapt the atmospheric pressure as a typical deep-sea bacterium as described previously (25).

**Table 2.**
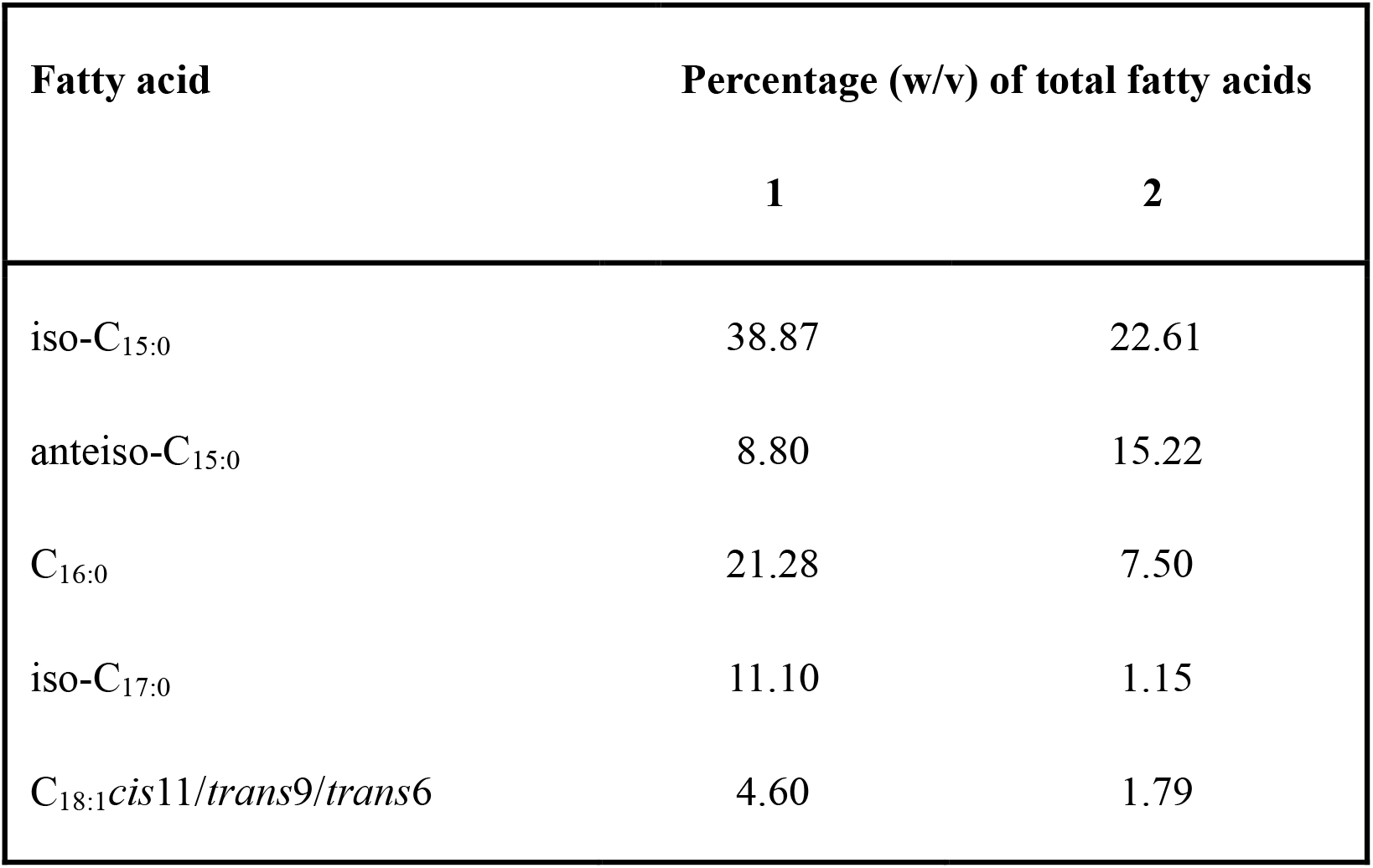
Comparison of the main fatty acids (%) of *P. cashew* SRB007^T^ with its closest relative *P. profundus* DSM 11384^T^. Strains: 1, *P. cashew* SRB007^T^ (all data from this study); 2, *P. profundus* DSM 11384^T^ (all data from this study).

### Dissimilatory sulfate reduction related genes existing in the genome of *P. cashew* SRB007

To gain more insights into sulfate reduction related characteristics of *P. cashew* SRB007, its whole genome was sequenced (Fig. S4). The genome size of *P. cashew* SRB007 was 3,909,950 bp with a DNA G+C content of 59.94%. The number of contig was 1, the total of N50 was 3,909,950 and the sequencing depth was 50.0×. Annotation of the genome of *P. cashew* SRB007 consisted of 3,499 coding sequences that included 68 RNA genes (9 rRNA genes, 55 tRNA genes and 4 other ncRNAs).

Notably, *P. cashew* SRB007 contained a complete gene cluster that composed of genes encoding some accessory factors and proteins closely related to dissimilatory sulfate reduction (Fig. 2A). Among them, *dsrAB*, *dsrC* together with *dsrMKJOP* encode all the necessary components of the Dsr complex required for sulfate reduction (26). And in this complex, DsrA and DsrB are annotated as different subunits of sulfite reductase; DsrC is annotated as sulfur relay protein; DsrK, DsrM and DsrP are annotated as menaquinone oxidoreductase; DsrJ is annotated as triheme cytochrome C; DsrO is annotated as 4Fe-4S ferredoxin iron-sulfur binding domain protein. The other three genes (*dsrD*, *dsrE* and *dsrN*) which are commonly present in *dsr* operons and encode proteins not directly involved in electron shuttling during sulfate reduction, were also identified in the genome of *P. cashew* SRB007. In combination of the discovery of other genes involved the sulfate reduction in the genome of *P. cashew* SRB007, an inferred pathway of dissimilatory sulfate reduction mediating reduction of sulfate to sulfide was shown in Figure 2B. In this pathway, SO_4_^2-^ was reduced to SO_3_^2-^ by a combination of Sat and AprAB; then SO_3_^2-^ was reduced to DsrC trisulfide by DsrAB and DsrC; the produced DsrC trisulfide was finally reduced to sulfide by the DsrMKJOP complex.

**FIG 2.**
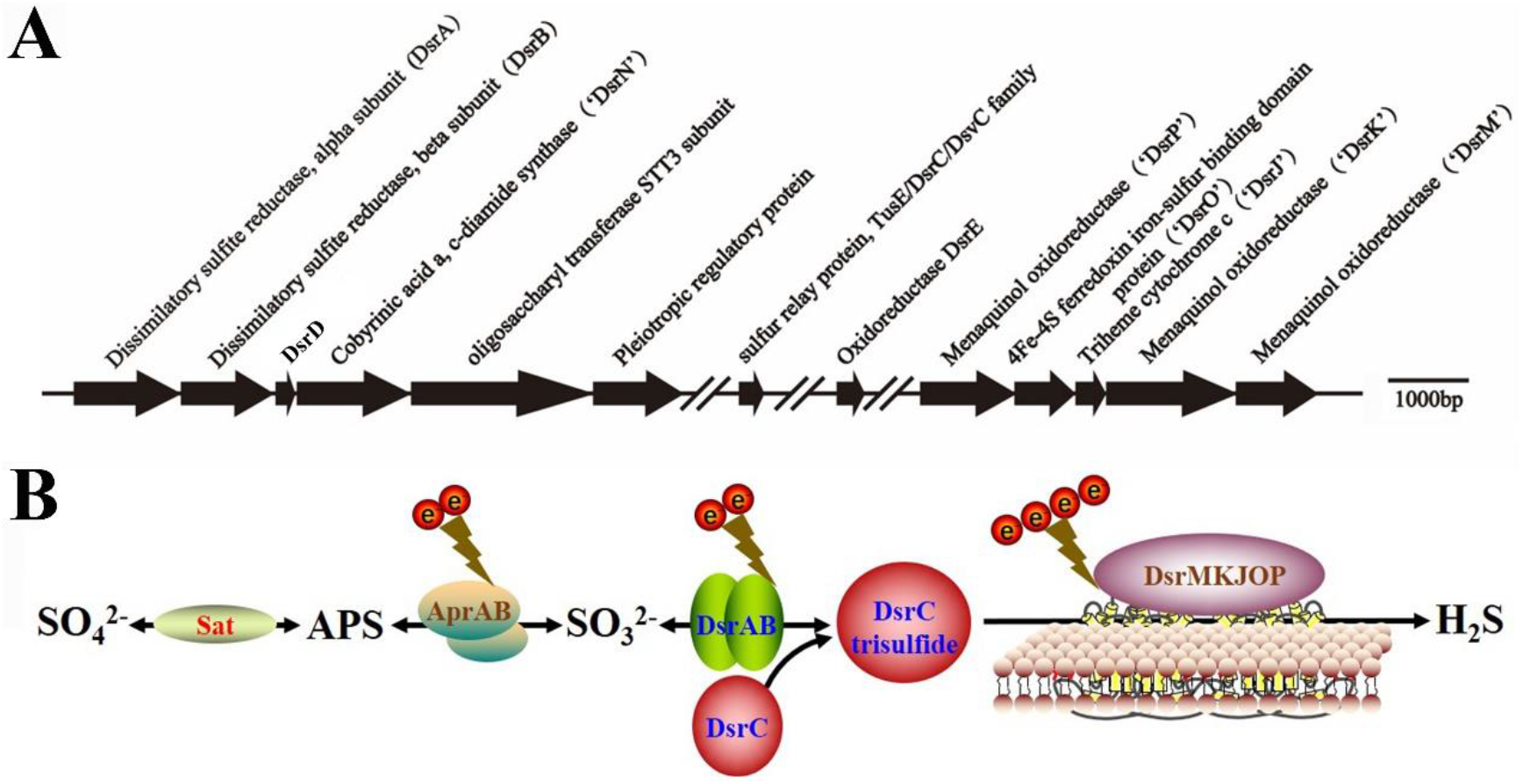
Genomic analysis of dissimilatory sulfate reduction of *P. cashew* SRB007. (A) The gene cluster containing the typical dissimilatory sulfate reductase operon and associated genes identified in the genome of *P. cashew* SRB007. (B) Proposed pathway of dissimilatory sulfate reduction of *P. cashew* SRB007. Sat, sulfuradenylyltransferase; APS, adenylyl sulfate; AprAB, adenylylsulfate reductase; DsrABC, reverse-type dissimilatory sulfite reductase; DsrMKJOP, sulfite reduction-associated complex.

### Dissimilatory sulfate reduction related genes contribute to the prominent capability of *P. cashew* SRB007 against different heavy metals

Given that *P. cashew* SRB007 is a typical sulfate-reducing bacterium, we next sought to explore its resistance to heavy metals and evaluate its potentials in the field of bioremediation as previously described (9, 10). With this, the growth status of *P. cashew* SRB007 was checked when challenged with different heavy metals including Fe^3+^, Co^2+^, Ni^2+^, Cu^2+^, Cd^2+^ and Hg^2+^, which are common metal ions existing in the deep-sea and the industrial waste water. The results showed that *P. cashew* SRB007 had an MIC (minimum inhibitory concentration) of 4.0 mM, 2.5 mM, 2.5 mM, 2.0 mM, 2.0 mM and 0.10 mM toward Fe^3+^, Co^2+^, Ni^2+^, Cu^2+^, Cd^2+^ and Hg^2+^, respectively (Fig. 3). Furthermore, the removal capabilities of *P. cashew* SRB007 toward the above tested heavy metals were checked. The results showed that the removal rate of Fe^3+^ was achieved 67.6% on the first day, then stabilized around 84.9% even extended the cultivation time to four days. Similarly, with the extension of culturing time, the removal rate of other heavy metals gradually increased and finally stabilized at 91.5%, 90.2%, 86.0%, 96.2% and 89.8% toward Co^2+^, Ni^2+^, Cu^2+^, Cd^2+^ and Hg^2+^,respectively, at the end of the fourth day (Fig. 4). It is noting that there were obvious precipitates formed in the bottom of the medium when *P. cashew* SRB007 was cultured with different metals, and the amount of precipitates increased with the length of incubation time. Given that *P. cashew* SRB007 is a typical SRB and has the potentials to form S^2-^, we proposed that these precipitates were metal sulfides. Indeed, these precipitates were further demonstrated to be Fe_2_S_3_, CoS, NiS, CuS, CdS and HgS by the energy dispersive spectrometry (EDS) analyses (Fig. 5), strongly indicating that dissimilatory sulfate reduction contributes to the resistance and removal capabilities of *P. cashew* SRB007 against different heavy metals. Taken together,*P. cashew* SRB007 had a prominent removal rate toward different heavy metals (Fe^3+^, Co^2+^, Ni^2+^, Cu^2+^, Cd^2+^ and Hg^2+^) via forming insoluble metal sulfides, indicating *P. cashew* SRB007 might be applied to the heavy metals treatment of sewage in the future.

**FIG 3.**
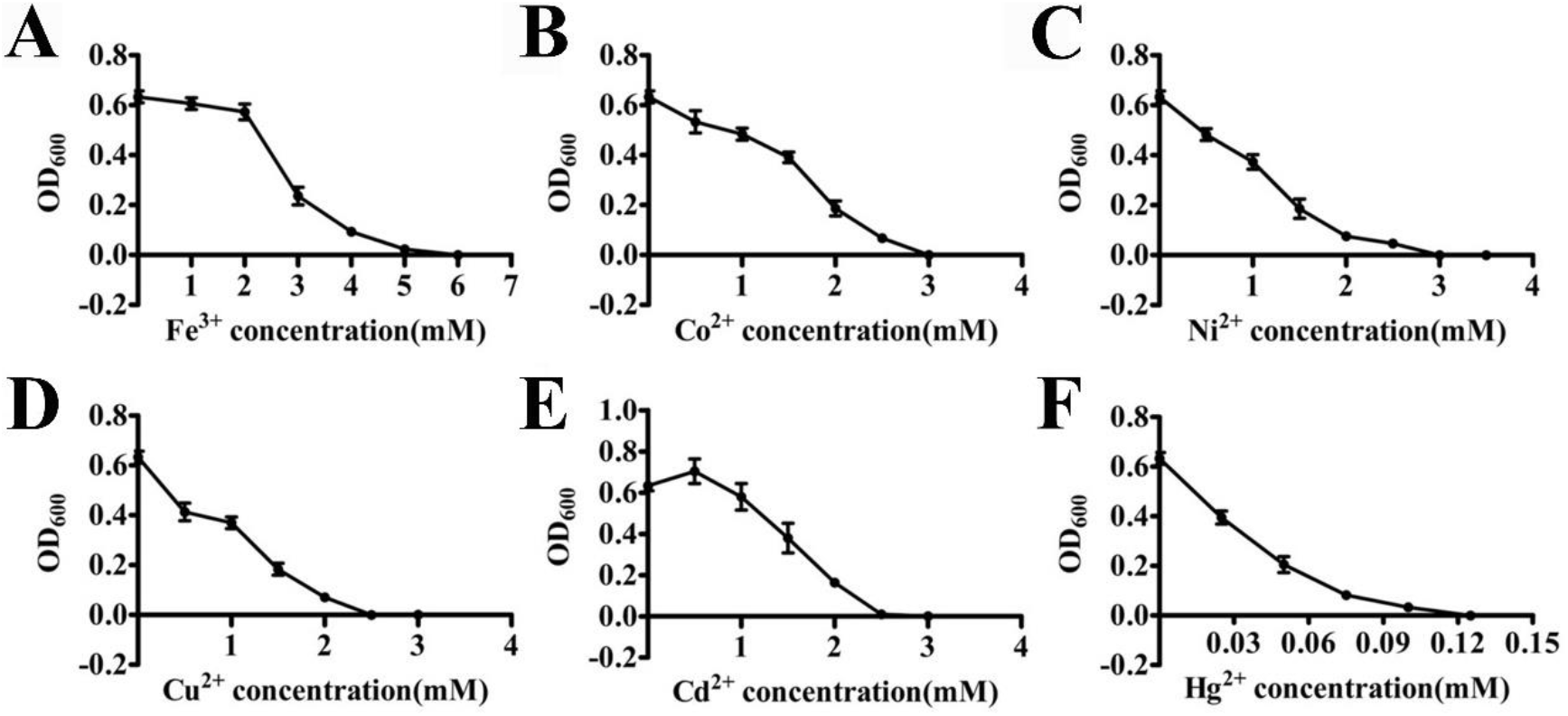
Determination of the minimum inhibitory concentration of Fe^3+^ (A), Co^2+^ (B), Ni^2+^ (C), Cu^2+^ (D), Cd^2+^ (E) and Hg^2+^ (F) against *P. cashew* SRB007.

**FIG 4.**
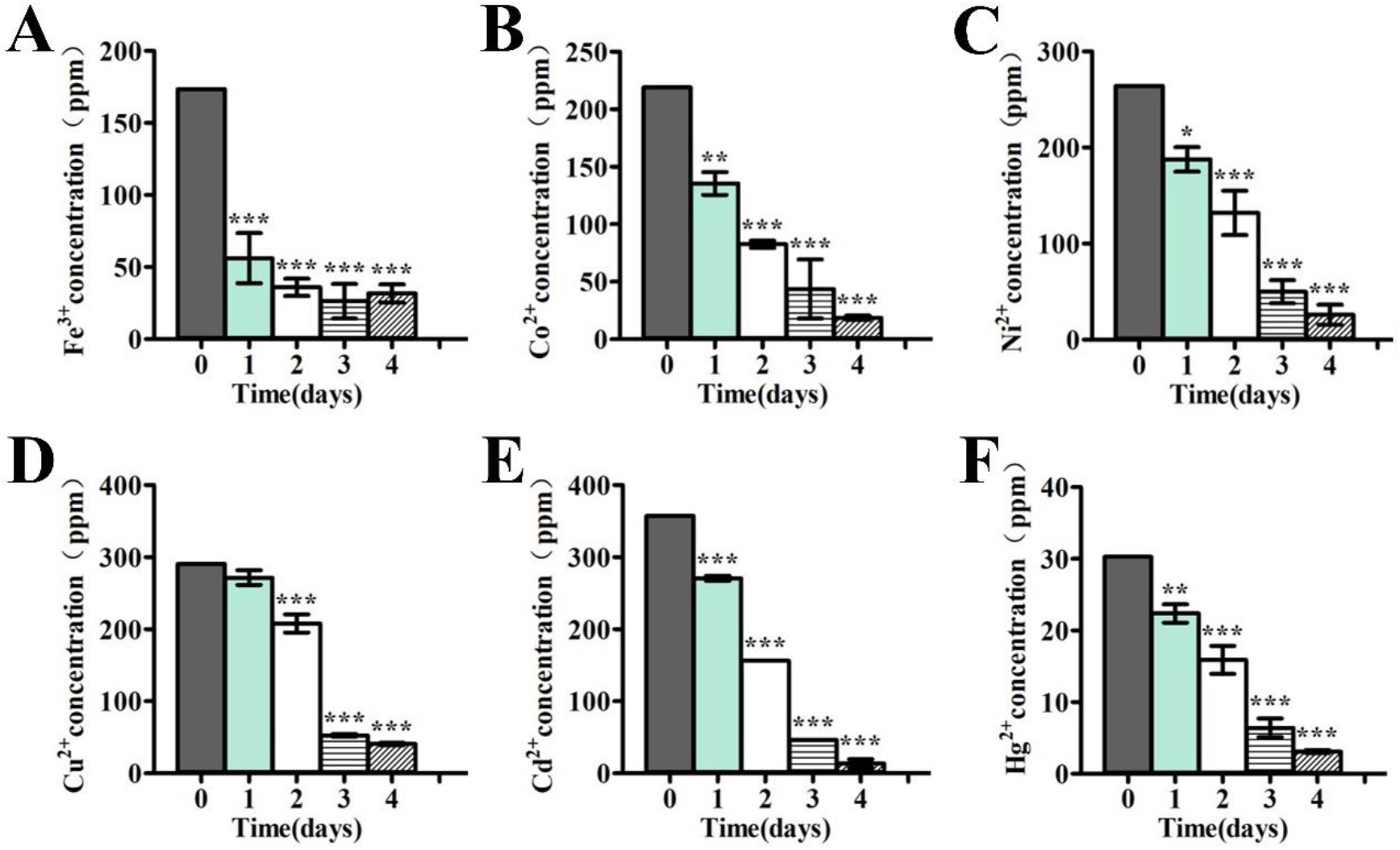
Mesurement of the removal efficiency of Fe^3+^ (A), Co^2+^ (B), Ni^2+^ (C), Cu^2+^ (D),Cd^2+^ (E) and Hg^2+^ (F) by *P. cashew* SRB007.

**FIG 5.**
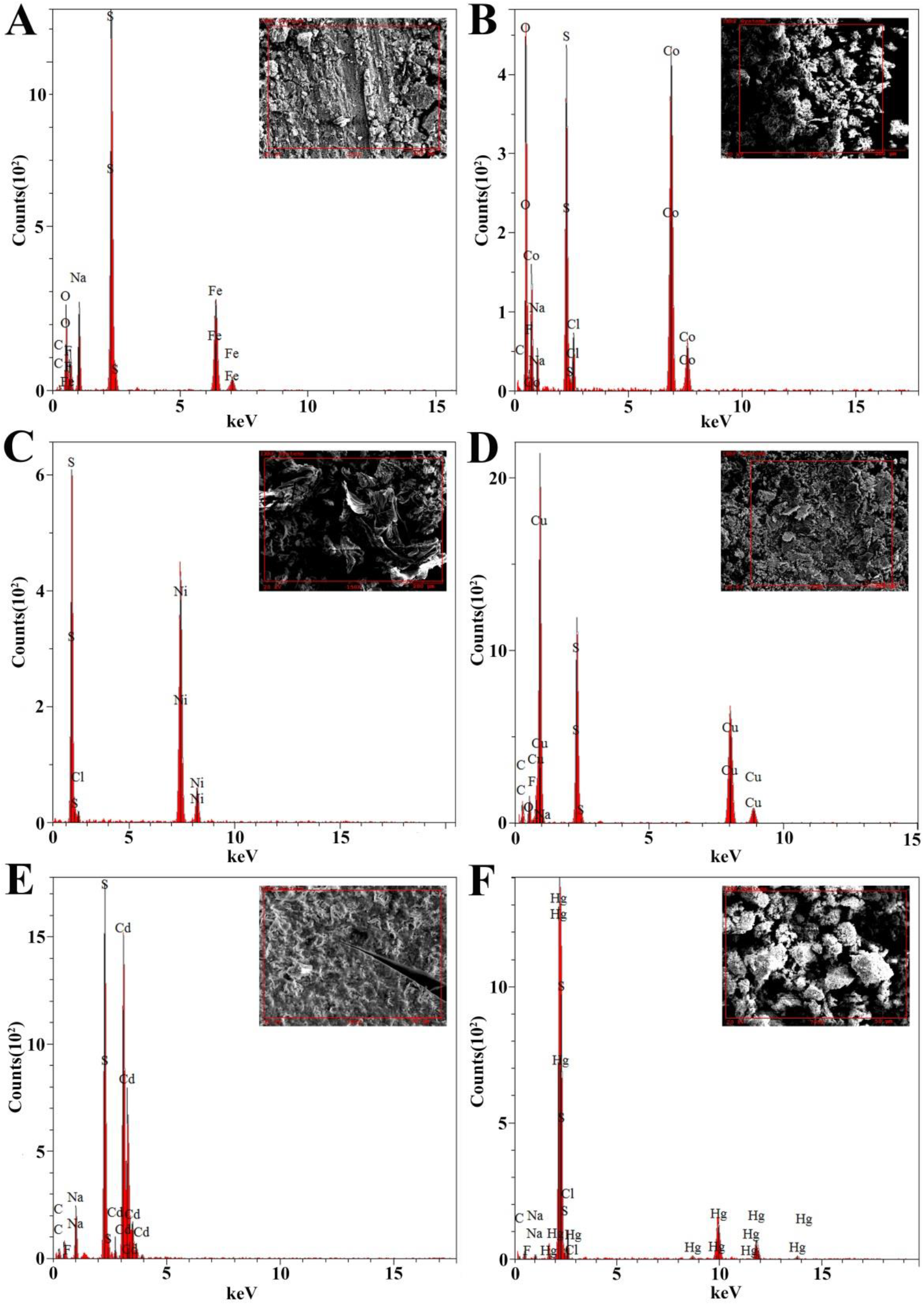
Energy dispersive spectrometry spectra of the precipitates (the inlet showing in each panel) formed by *P. cashew* SRB007 when cultured in the medium containing different concentrations of Fe^3+^ (A), Co^2+^ (B), Ni^2+^ (C), Cu^2+^ (D), Cd^2+^ (E) and Hg^2+^(F), respectively.

To confirm the participation of the dissimilatory sulfate reduction associated genes in the course of coping with different heavy metals by *P. cashew* SRB007, the expressions of several important genes within the gene cluster mentioned in Figure 2A were checked by real-time quantitative PCR. Clearly, the expression of *dsrABDNCE* was significantly up-regulated from ~6- to ~100- fold when challenged with Cd^2+^ and Hg^2+^. However, the expression of *dsrABE* was only up-regulated from ~2- to ~5- fold when challenged with Fe^3+^, Co^2+^, Ni^2+^ and Cu^2+^ (Fig. 6). Given that Cd^2+^ and Hg^2+^ are high toxicity metal ions and Fe^3+^, Co^2+^, Ni^2+^ and Cu^2+^ are essential elements to life, it is reasonable to see the results that the expression of *dsrABDNCE* was much higher when challenged with Cd^2+^ and Hg^2+^ comparing with other low toxicity metal ions like Fe^3+^, Co^2+^, Ni^2+^ and Cu^2+^.

**FIG 6.**
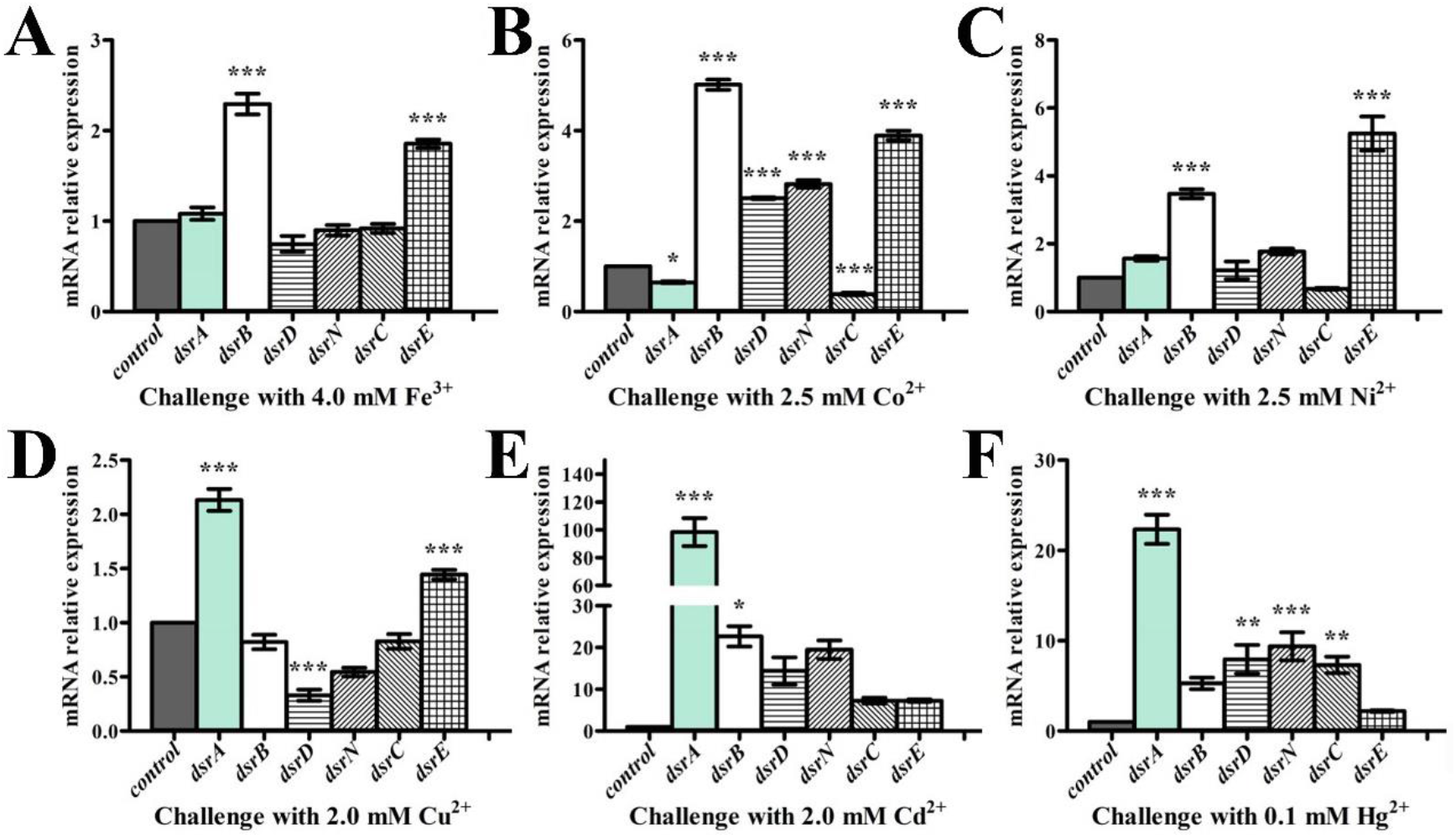
qRT-PCR analysis of the expression of genes within *dsr*-operon of *P. cashew* SRB007 challenged with 4.0 mM Fe^3+^ (A), 2.5 mM Co^2+^ (B) 2.5 mM Ni^2+^ (C) 2.0 mM Cu^2+^ (D), 2.0 mM Cd^2+^ (E) and 0.1 mM Hg^2+^ (F), respectively.

### Proposed life style of *P. cashew* SRB007

In combination of the genomic and physiological traits of *P. cashew* SRB007, a proposed life style was shown in Figure 7. First, there were corresponding genes for utilizing acetate and lactate in the genome of *P. cashew* SRB007, which was consistent with our results shown in Table 1 that acetate and lactate could be used as electron donors and carbon sources of this bacterium (Fig. 7). In addition, the genes in charge of glycolysis and oxidative pentose phosphate biosynthesis were also discovered in the bacterial genome (Fig. 7). Overall, *P. cashew* SRB007 could behave a heterotrophic life via consuming organic matter like acetate and glucose to produce acetyl-CoA and generate energy by tricarboxylic acid cycle. On the other hand, many genes responsible for autotrophic were also found in the genome of *P. cashew* SRB007. Such as, nearly all of the genes involved in the nitrogen fixation process including *nifHDK* encoding molybdenum-iron nitrogenase, *nifBENSU* encoding nitrogen-fixing assembly proteins and *nifA* encoding transcriptional regulator proteins (Table S3) (27); genes encoding the complete components of NiFe hydrogenase, Ech hydrogenase, malate dehydrogenase, formate dehydrogenase, succinate dehydrogenase and alcohol dehydrogenase (Table S3); genes encoding anaerobic carbon-monoxide dehydrogenase, which catalyzed the oxidation of carbon monoxide to carbon dioxide or the reverse reaction. Together,*P. cashew* SRB007 could behave autotrophic life via utilizing N_2_, H_2_, CO and CO_2_ to produce energy. Finally,*P. cashew* SRB007 also possesses the sodium translocating NADH: Ferredoxin oxidoreductase (RNF complex), the proton translocating NADH: Quinone oxidoreductase (complex I) and cytochrome *bd* ubiquinol oxidase (cydAB). Both the complex I and cytochrome *bd* ubiquinol oxidase interact with the menaquinone pool, which can form a simple electron transport chain to generate energy (28). All the above results indicate that *P. cashew* SRB007 could generate energy via multiple pathways, which provides enough energy for bacterial sulfur cycle and stress tolerance.

**FIG 7.**
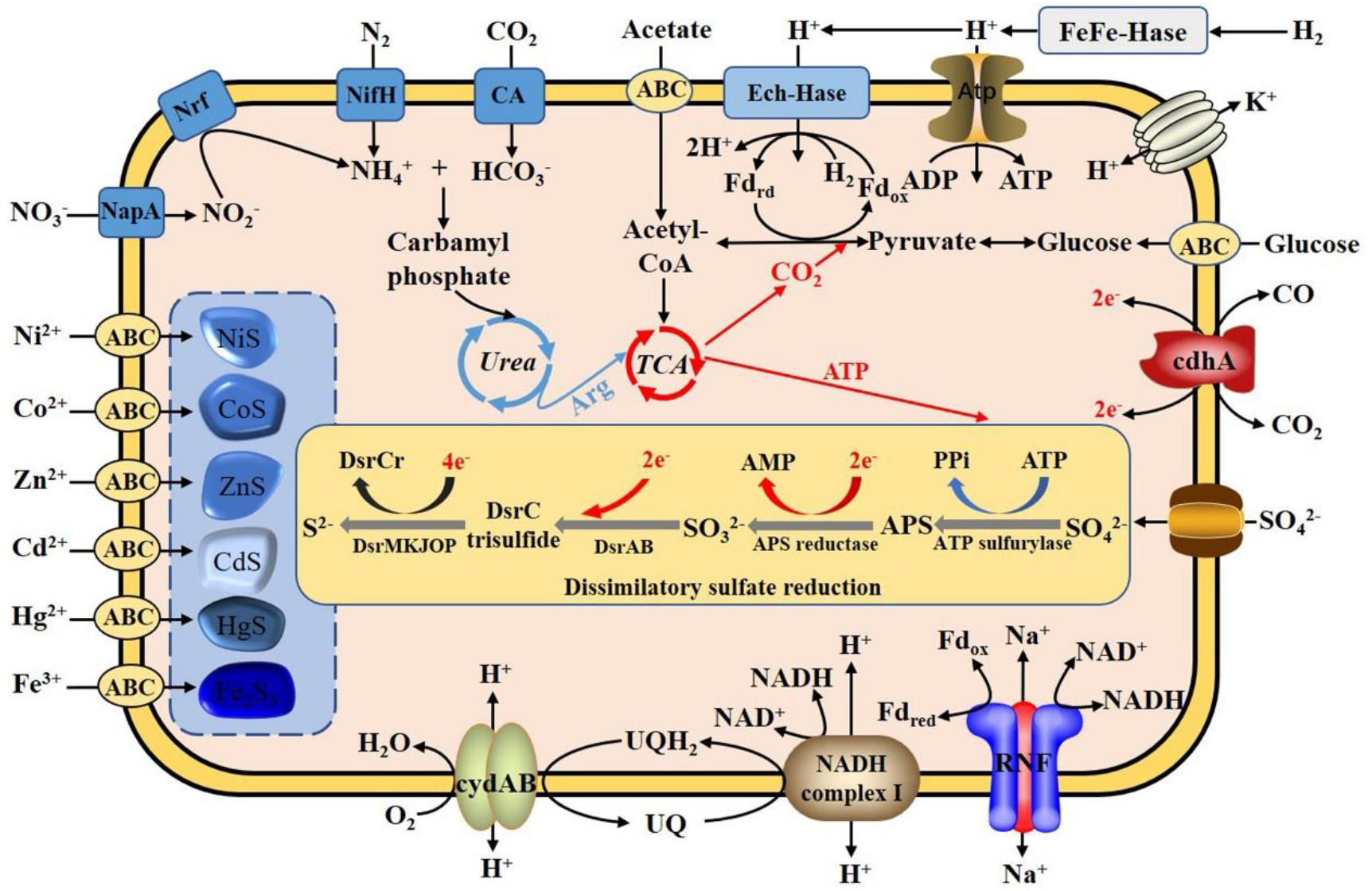
Proposed lifestyle of *P. cashew* SRB007. Abbreviations: TCA, tricarboxylic acid cycle; Urea, urea cycle; FeFe-Hase, FeFe-hydrogenase; Atp, ATP synthase; ATP, 5′-Adenylate triphosphate; ADP, adenosine diphosphate; AMP, adenosine monophosphate; Ech-Hase, energy-conserving membrane-bound hydrogenase; CA, carbonic anhydrase; NifH, nitrogenase iron protein; Nrf, cytochrome c nitrite reductase; NapA, nitrate reductase catalytic subunit; cydAB, cytochrome bd-I ubiquinol oxidase subunit; cdhA, CO dehydrogenase/acetyl-CoA synthase complex subunit epsilon; UQ, ubiquinone; UQH2, Cytochrome C reductase; RNF, the sodium translocating NADH: Ferredoxin oxidoreductase; DsrAB, dissimilatory sulfite reductases; PPi, pyrophosphoricacid.

## DISCUSSION

Sulfur is a key element in the nature, whose transformation and status are critically dependent upon microbial activities. The sulfur cycle in marine sediments is primarily driven by the dissimilatory sulfate reduction to sulfide by the anaerobic SRB (1). There have been extensive studies of sedimentary sulfur cycle across the global ocean and focus on geological microscopic transformations of the two end-members sulfate and sulfide (29). The deep-sea cold seep is a very special environment where is abundant methane and sulfate and estimates of quantities in the sediments of cold seep suggest that SRB account for approximately 5-25% of the microbial biomass in the surface of sulfate-rich zones and may reach higher up to approximately 30-35% in the sulfate-methane transition zone (30). Therefore, the deep-sea cold seep is one of the best locations to study the sulfur cycle mediated by SRB. However, because of the difficulty of sample collection and absence of pure SRB cultures from the deep-sea cold seep, it is utmost precious to obtain the typical SRB to explore the unknown mechanisms about sulfur cycle happened in this special environment.

In the present study, a novel sulfate-reducing bacterium designated *P. cashew* SRB007 was isolated and purified from the deep-sea cold seep and proposed to represent a novel species in the genus of *Pseudodesulfovibrio* (Fig. 1). Till to date, only five species were reported in the genus of *Pseudodesulfovibrio* and four of them were isolated from marine sediments, including *P. profundus* (16), *P. portus* (17), *P. piezophilus* (18) and *P. indicus* (15). *P. cashew* SRB007 reported in this study is the first species isolated from the deep-sea cold seep of the genus of *Pseudodesulfovibrio. P. cashew* SRB007 can grow well under the conditions of 16 to 45 °C, 0-100 g/L NaCl and utilize fumarate, acetate, methanol, ethanol, formate, lactate, succinate, malate as electron donors and sulfate, sulfite, thiosulfate, nitrate, nitrite as electron acceptors (Table 1). Thus it can be seen that *P. cashew* SRB007 possesses a wide growth condition and a broad range of substrates involving in sulfur and nitrogen cycles.

Given the isolation location of *P. cashew* SRB007, it is reasonable to speculate that this bacterium should have a strong capability to resist different heavy metals. As respected, *P. cashew* SRB007 could resist and remove high concentrations of Fe^3+^, Co^2+^, Ni^2+^, Cu^2+^, Cd^2+^ and Hg^2+^ (Figs. 4 and 5). Previous research has reported that heavy metal removal and sulfate reduction correlated with each other (31). As *P. cashew* SRB007 is a sulfate-reducing bacterium, it is logical to infer that this bacterium removes heavy metals through sulfate reduction to form metal sulfide precipitation. Indeed, the expression of the dissimilatory sulfite reduction genes (*dsrABCDEMNKJOP*) of *P. cashew* SRB007 was demonstrated to correlate with its heavy metals resistance and removal capabilities (Figs. 2 and 6). Overall,*P. cashew* SRB007 is a novel deep-sea sulfate-reduction bacterium that growing well in the atmospheric pressure and possessing a strong capability for removing many harmful heavy metals.

It is noting that many metal-polluted wastewater, such as acid mine wastewater and industrial metallurgic wastewater, contains high concentrations of sulfate and toxic heavy metals, which is pretty similar to that of deep-sea and poses a serious threat to the environment when discharged untreated (32,33). Based on the understanding of the detoxification mechanism used by SRB to reduce the toxicity of heavy metals, it is possible to develop efficient and environment friendly metal repair technology. Undoubtedly,*P. cashew* SRB007 is a good candidate to develop the corresponding bioremediation product as shown in Fig. S6 in the future, given its strong environmental adaptative and heavy metal resistant capabilities.

Furthermore, sulfur cycle is tightly interwoven with other important element cycles (carbon, nitrogen, manganese and iron) in marine sediments (2). Till to date, most of the mechanisms about sulfur cycle and its coupling with other elements mediated by SRB are uncovered. Given that *P. cashew* SRB007 is a typical deep-sea cold seep sulfate-reducing bacterium and possesses both heterotrophic and autotrophic lifestyles by using many organic and inorganic matter as the energy sources (Fig. 7), it will be a great interest to deeply explore the mechanisms of sulfur cycle linking with other elements of this bacterium in the future.

## MATERIALS AND METHODS

### Bacterial strains and culture conditions

The samples were collected by *RV KEXUE* from the cold seep in the South China Sea (E 119°17′07.322″ N 22°06′58.598″) as described previously (3, 34). And the sediment samples were cultured at 28 °C for one month in an anaerobic enrichment medium containing (per litre of seawater): 1 g NH_4_Cl, 1 g NaHCO_3_, 1 g CH_3_COONa, 0.5 g KH_2_PO_4_, 0.2 g MgSO_4_.7H_2_O, 1 g peptone, 1 g yeast extract, 0.7 g cysteine hydrochloride, 1 mL 0.1% (w/v) resazurin (the pH was adjusted to 7.0) and the medium was prepared anaerobically as previously described (35). The cultures were purified by repeated use of the Hungate roll-tube method. Single colonies were picked by sterilized bamboo skewers and then cultured in the same medium. The process of isolation was repeated several times until the isolates were deemed to be axenic. The purity of the isolate was confirmed routinely by transmission electron microscopy (TEM) and by repeated partial sequencing of the 16S rRNA gene. Then the single colony was transferred to a new medium (D195c) containing (per litre of seawater): 1.0 g yeast extract, 2.0 g peptone, 2.2 g sodium lactate, 2.0 g Na_2_SO_4_, 3.3 g PIPES, 1.0 g cysteine hydrochloride, 1 mL 0.1% (w/v) resazurin; the pH of the medium was adjusted to 7.0 with NaOH. The cultures were maintained at 30 °C in an anaerobic chamber (Longyue, China) under a gas mixture of 90% N_2_, 5% CO_2_ and 5% H_2_.

### Transmission electron microscopy (TEM) observation

To observe the morphological characteristics of *P. cashew* SRB007, the cells were examined using TEM with a JEOL JEM 12000 EX (equipped with a field emission gun) at 100 kV. The cell suspension of *P. cashew* SRB007 was washed with Milli-Q water and centrifuged at 4,000 × *g* for 5 min. Subsequently, the sample was taken by immersing copper grids coated with a carbon film for 20 min in the bacterial suspensions and washed for 5 min in distilled water and dried for three hours at room temperature (36).

### Genomic sequencing and analysis

To obtain the whole genome of *P. cashew* SRB007, total chromosomal DNA of this bacterium was extracted. The DNA library was prepared, using the Ligation Sequencing Kit (SQK-LSK109) and sequenced, using a FLO-MIN106 vR9.4 flow-cell for 48 h on MinKNOWN software v1.4.2 (Oxford Nanopore Technologies (ONT), United Kingdom). Whole-genome sequence determination was carried out with the Oxford Nanopore MinION (Oxford, United Kingdom) and Illumina MiSeq sequencing platform (San Diego, CA) and a hybrid approach was further utilized for genome assembly using reads from both platforms. Base-calling was performed via Albacore software v2.1.10 (Oxford Nanopore Technologies). Nanopore reads were processed with Poretools toolkit for the purposes of quality control and downstream analysis (37) and the filtered reads were assembled by Canu version 1.8 (38). The whole genome was finally assembled into a single contig and was manually circularized by deleting an overlapping end. Based on the whole genome of *P. cashew* SRB007, full-length 16S rRNA,*dsrABCDE* and other genes related to sulfate reduction were obtained and applied to different analyses.

### Phylogenetic analysis

Sequences of 16S rRNA and *dsrAB* of *P. cashew* SRB007 and other related taxa used for phylogenetic analysis were all obtained from NCBI GenBank. Phylogenetic analysis was performed using the software MEGA version 6.0 (39). The phylogenetic tree was constructed by the neighbor-joining algorithm (40), maximum likelihood (41) and minimum-evolution methods (42). The numbers above or below the branches were bootstrap values based on 1,000 replicates.

### Physiological and chemotaxonomic assays of *P. cashew* SRB007

Morphological characteristics and purity of *P. cashew* SRB007 were observed by TEM. Growth assays were performed at atmospheric pressure, using Hungate tubes containing basal medium (3.3 g PIPES, 1.0 g cysteine hydrochloride, 1 mL 0.1% (w/v) resazurin; the pH of the medium was adjusted to 7.0 with NaOH) and different electron donors at 20 mM (acetate, fumarate, formate, pyruvate, lactate, malate, methanol, fructose, propionate, butyrate, succinate, glycine, ethanol). Elemental sulfur (1%, w/v), sulfate (20 mM), sulfite (20 mM), thiosulfate (20 mM), nitrate (10 mM) and nitrite (10 mM) were tested as terminal electron acceptors. The temperature, pH and NaCl concentration ranges for the growth of *P. cashew* SRB007 were determined in duplicate experiments using autotrophic medium supplemented with lactate (20 mM) as electron donor and sulfate (20 mM) as previously described (43). Temperatures for growth were tested between 4 and 80 °C. The pH range for growth was tested from pH 3.0 to pH 11.0 (at 30 °C) with increments of 0.5 pH units. Salt resistance was determined by directly weighing NaCl (0-100 g L^-1^) into the Hungate tubes before packaging the autotrophic medium.

### Determination of minimal inhibitory concentrations (MIC) of *P. cashew* SRB007 challenged with different heavy metal ions

To take the determination of Fe^3+^ (Iron III) MIC against *P. cashew* SRB007 as an example, 10 μL of the cultured bacterial solution was inoculated in 5 mL D195c anaerobic medium containing 0, 1, 2, 3, 4, 5, 6, 7, 8, 9 or 10 mM FeCl_3_, respectively. The cultures were incubated at 30 °C in the anaerobic chamber for 48 h and then determined by spectrophotometry at 600 nm. The MIC value is the lowest concentration of heavy metal ions at which growth in the D195c anaerobic medium was inhibited (44). And the determination of other heavy metal ions Co^2+^, Ni^2+^, Cu^2+^, Cd^2+^ and Hg^2+^ against *P. cashew* SRB007 was performed as mentioned above.

### Heavy metal removal assay and qualitative energy dispersive spectrometry (EDS) analysis

To take the determination of Hg^2+^ removal rate of *P. cashew* SRB007 as an example, *P. cashew* SRB007 was incubated at 30 °C in D195c anaerobic medium supplemented with 0.1 mM HgCl_2_ to OD_600_ value of 1.0. The supernatant of culture was collected by centrifugation (12,000 × *g*, 5 min). After this, the supernatant was thoroughly digested with perchloric acid and nitric acid and diluted with Milli-Q water for Hg^2+^ concentration detection. The dissolved Hg^2+^ concentrations were measured with an inductively coupled plasma source mass spectrometer (Optima 7300 DV, PerkinElmer). And the determination of the removal rate of other heavy metal ions (Fe^3+^, Co^2+^, Ni^2+^, Cu^2+^, Cd^2+^) against *P. cashew* SRB007 was performed as mentioned above. In addition, the precipitation was washed with Milli-Q water for three times and then performed ultrasonic decomposition for 30 min. The sample was collected by centrifugation (12,000 × *g*, 10 min) and washed wth Milli-Q water for three times. Finally, the sample was dried in an oven for 3 h at 80 °C and then thin coated by Au for EDS analysis using a model 550i (IXRF SYSTEMS, America).

### RNA extraction, reverse transcription and quantitative real-time PCR (qRT-PCR)

For qRT-PCR, cells of *P. cashew* SRB007 challenged with different heavy metal ions (4.0 mM Fe^3+^, 2.5 mM Co^2+^, 2.5 mM Ni^2+^, 2.0 mM Cu^2+^, 2.0 mM Cd^2+^ and 0.1 mM Hg^2+^) were grown in D195c anaerobic medium at 30 °C for 2 days, then 2 mL of these cells were harvested and centrifuged at 12,000 ×*g* for 5 min, respectively. Total RNAs were extracted using the Trizol reagent (Solarbio, China) and the RNA concentration was measured using Qubit^®^ RNA Assay Kit in Qubit® 2.0 Fluorometer (Life Technologies, CA, USA). Then RNA was reverse transcribed into cDNA and the transcriptional levels of different genes were determined by qRT-PCR using SybrGreen Premix Low rox (MDbio, China) and the QuantStudio™ 6 Flex (Thermo Fisher Scientific, America). 16S rRNA was used as an internal reference and the *merF* gene expression was calculated using the 2^−ΔΔCt^ method, with each transcript signal normalized to 16S rRNA (45, 46). Transcript signals for each treatment were compared to the transcript signals of the control group. The transcription levels of each *dsr* gene were normalized to the 16S rRNA gene. Specific primers for *dsrA*, *dsrB*, *dsrD*, *dsrN*, *dsrC*, *dsrE* and 16S rRNA of *P. cashew* SRB007 were designed using Primer 5.0 (Table 3). All qRT-PCR runs were performed in three biological and three technical replicates.

**Table 3.**
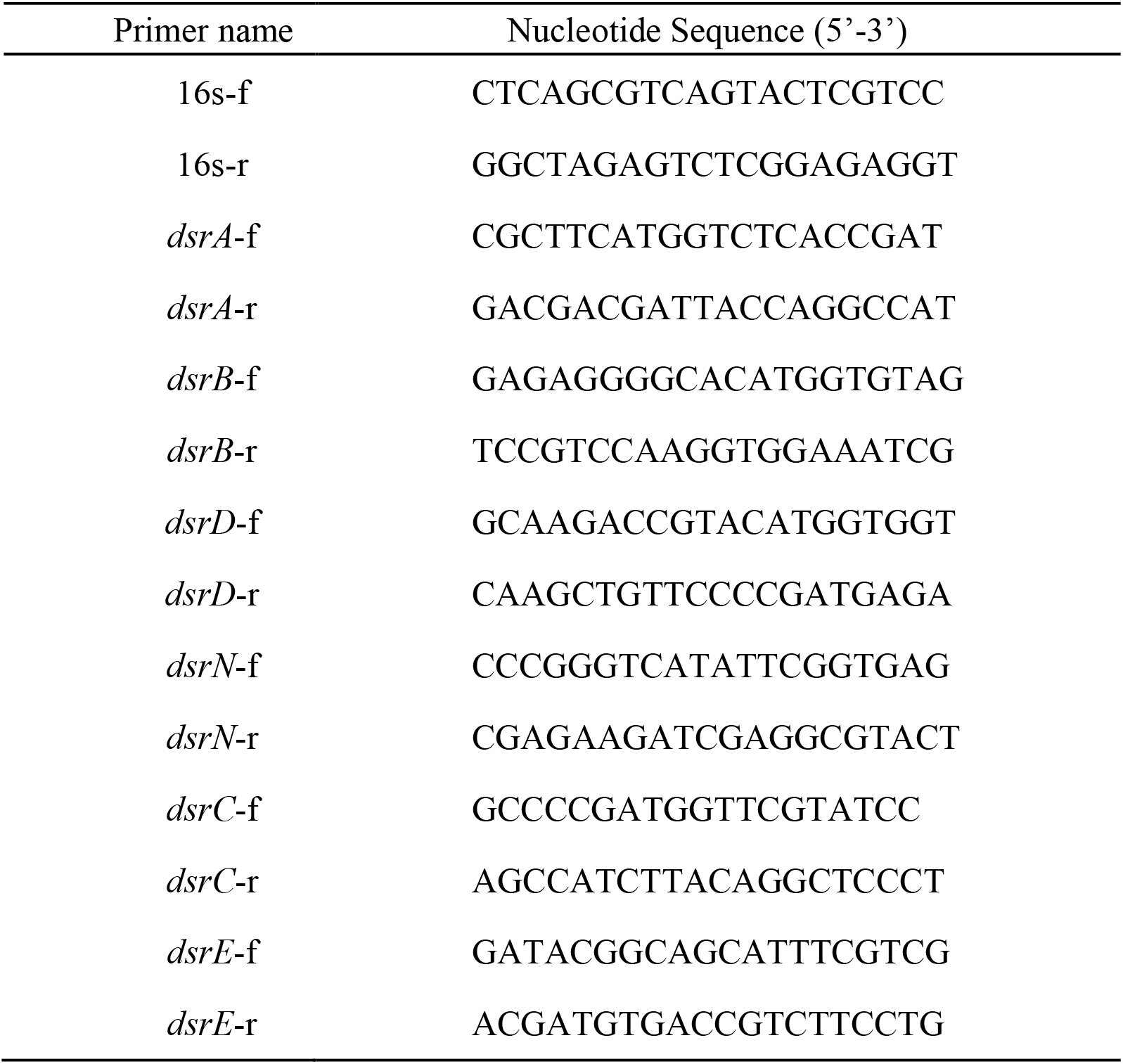
Primers used for qRT-PCR in this study.

### Statistical Analysis

Statistical analysis in this study was performed using GraphPad Prism 5, tests for significance of the differences among groups were subjected to one-way analysis of variance (one-way ANOVA) and multiple comparisons. A statistically significance was defined in our study by *P*< 0.05 (indicated by* in all figures), *P*< 0.01 (indicated by ** in all figures) or *P*< 0.001 (indicated by *\*\*\** in all figures).

### Data deposit

The complete genome sequence of *P. cashew* SRB007 has been deposited at GenBank under the accession number CP046400. The NCBI GenBank accession number for the 16S rRNA gene sequence of *P. cashew* SRB007 is AF418172.

### Description of *Pseudodesulfovibrio cashew* sp. nov

*Pseudodesulfovibrio cashew* (ca’sh.ew N.L. n. cashew nut, referring to the character is similar to a cashew nut).

Cells of strain SRB007^T^ are Gram-stain-negative, strictly anaerobic, cashew-shaped, 1.0-2.5 μm in length and 0.3-0.7 μm in width, motile by peritrichous flagella. The temperature range for growth is 16-45 °C with an optimum at 30 °C. Growing at pH values of 5.5-8.5 (optimum, pH 7.0) and at NaCl of 0-100 g/L (optimum, 50 g/L). Fumarate, acetate, methanol, ethanol, formate, lactate, succinate and malate are oxidized with sulfate reduction. Sulfate, sulfite, thiosulfate, nitrate and nitrite serve as electron acceptors. The major polar lipids are phosphatidylethanolamine, diphosphatidylglycerol, phosphatidylglycerol, unidentified glycolipid and unknown aminoglycolipids. Containing significant proportions (>10%) of the cellular fatty iso-C_15:0_, C_16:0_, iso-C_17:0_.

The type strain, SRB007^T^ (=KCTC 15990^T^=MCCC 1K04423^T^), was isolated from deep-sea sediments of cold seep, P.R. China. The DNA G+C content of the type strain is 59.94%.

## ACKNOWLEDGEMENTS

This work was funded by the the Major Research Plan of the National Natural Science Foundation (Grant No. 92051107), China Ocean Mineral Resources R&D Association Grant (Grant No. DY135-B2-14), Strategic Priority Research Program of the Chinese Academy of Sciences (Grant No. XDA22050301), National Key R and D Program of China (Grant No. 2018YFC0310800), the Taishan Young Scholar Program of Shandong Province (tsqn20161051), and Qingdao Innovation Leadership Program (Grant No. 18-1-2-7-zhc) for Chaomin Sun.

